# Pyruvate kinase deficiency links metabolic perturbations to neurodegeneration and axonal protection

**DOI:** 10.1101/2025.04.04.647282

**Authors:** Thomas J. Waller, Catherine A. Collins, Monica Dus

## Abstract

Neurons rely on tightly regulated metabolic networks to sustain their high-energy demands, particularly through the coupling of glycolysis and oxidative phosphorylation. Here, we investigate the role of pyruvate kinase (PyK), a key glycolytic enzyme, in maintaining axonal and synaptic integrity in the *Drosophila melanogaster* neuromuscular system. Using genetic deficiencies in PyK, we show that disrupting glycolysis induces progressive synaptic and axonal degeneration and severe locomotor deficits. These effects require the conserved dual leucine zipper kinase (DLK), Jun N-terminal kinase (JNK), and activator protein 1 (AP-1) Fos transcription factor axonal damage signaling pathway and the SARM1 NADase enzyme, a key driver of axonal degeneration. As both DLK and SARM1 regulate degeneration of injured axons (Wallerian degeneration), we probed the effect of PyK loss on this process. Consistent with the idea that metabolic shifts may influence neuronal resilience in context-dependent ways, we find that *pyk* knockdown delays Wallerian degeneration following nerve injury, suggesting that reducing glycolytic flux can promote axon survival under stress conditions. This protective effect is partially blocked by DLK knockdown and fully abolished by SARM1 overexpression. Together, our findings help bridge metabolism and neurodegenerative signaling by demonstrating that glycolytic perturbations causally activate stress response pathways that dictate the balance between protection and degeneration depending on the system’s state. These results provide a mechanistic framework for understanding metabolic contributions to neurodegeneration and highlight the potential of metabolism as a target for therapeutic strategies.

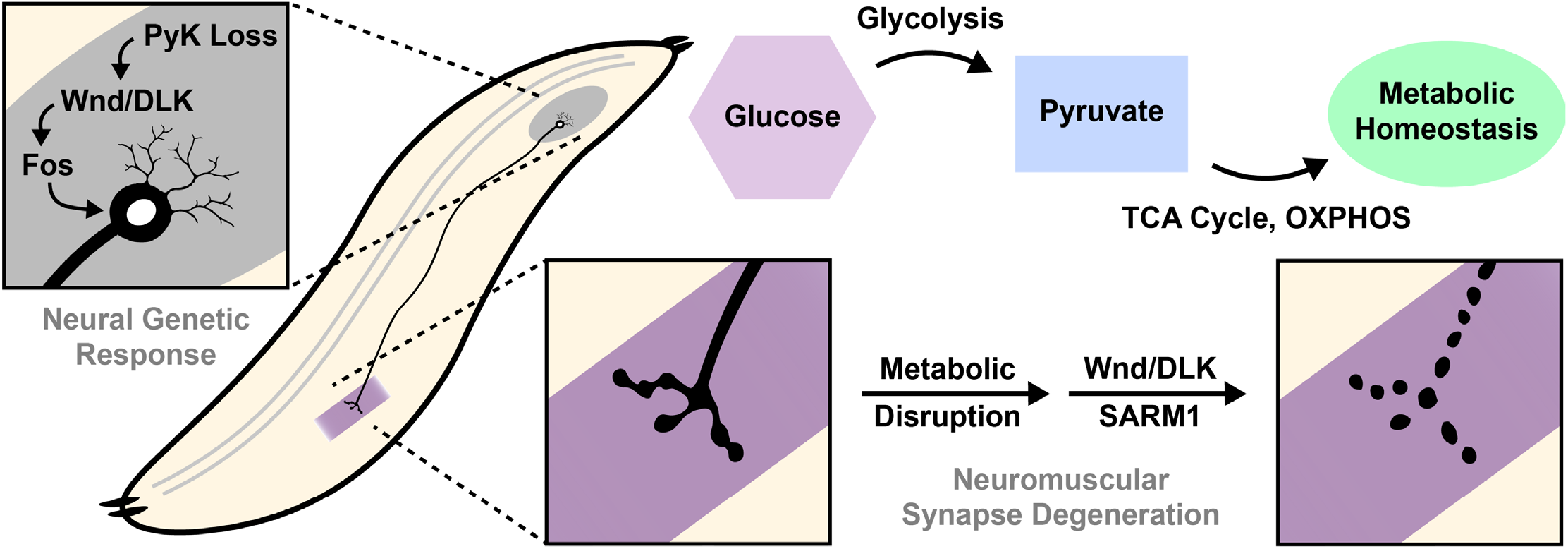

## Introduction

The nervous system’s ability to sustain rapid signaling relies on a continuous energy supply. Neurons must generate ATP constantly to maintain and restore the electrochemical gradients necessary for action potential propagation and neurotransmission. This energy is derived from a well-coordinated network of metabolic pathways, particularly those involving central carbon metabolism—glycolysis, the citric acid cycle, and oxidative phosphorylation—that provide fuel and the necessary biosynthetic precursors and redox equivalents for cellular function [1,2]. Motor neurons, in particular, depend heavily on these metabolic pathways to support their long axonal projections and high synaptic activity [3–5]. Perturbations in energy metabolism have been observed in both peripheral and central neurodegenerative disorders, although critical questions remain on whether these changes are the cause or consequence of neurodegeneration [6–10]. Further, not much is known on how metabolic perturbations interact with conserved, canonical pathways of neurodegeneration and regeneration. These include the mitogen-activated protein kinase (MAPK) c-Jun N-terminal kinase (JNK) and the upstream MAP3K dual leucine zipper kinase (DLK), which are activated by stress and axonal injury [11–19]. Further connections between metabolism and axonal integrity were recently highlighted by the discovery that one of the key mediators of axonal degradation, sterile alpha and TIR motif containing 1 (SARM1)—which drives this process through nicotinamide adenine dinucleotide (NAD) degradation—is regulated by levels of NAD and its precursor nicotinamide mononucleotide (NMN), making SARM1 both an affector and an effector of metabolic disruption [20–25]. The signaling mechanisms of these pathways are central to the development of biomarkers and therapeutic targets for central and peripheral nervous system diseases. As such, understanding their crosstalk with metabolic pathways is a question of urgent conceptual and clinical relevance.

Besides the intrinsic challenges of studying metabolism, which is inherently dynamic and in constant flux, another major difficulty has been the need for more in vivo systems that allow us to model metabolic perturbations, observe their effects on axonal integrity, and track the engagement of neurodegenerative signaling pathways in a physiologically relevant context. While in vitro studies have provided insights into metabolic perturbations, they do not always capture how these changes impact neuronal integrity within an intact physiological system, though technical advances continue to close this gap [26]. In vivo studies, on the other hand, often focus on identifying genetic and protein signaling neurodegenerative pathways without directly characterizing causal relationships between specific metabolic perturbations and these mechanisms [27].

Here, we bypassed these challenges by leveraging the unique advantages of the *Drosophila melanogaster* neuromuscular junction (NMJ), a model that has been instrumental to uncovering fundamental principles of neuronal communication, synaptic maintenance, and degeneration [28–32]. For decades, studies in the fly NMJ have revealed highly conserved mechanisms of neurodegenerative signaling, axonal transport, and metabolic regulation, bridging basic research with preclinical studies [33,34].

In this study, we took advantage of the NMJ’s well-characterized anatomy, regulatory mechanisms, and powerful genetic tools to dissect the interplay between metabolic disruption and neurodegenerative signaling. By doing so, we specifically aimed to define how metabolic perturbations observed in clinical settings impact axonal integrity and animal behavior in vivo, how they interact with canonical neurodegenerative pathways, and how the outcomes of this crosstalk control axon degeneration and synapse loss.

To model the disruption of metabolic homeostasis observed in human and rodent models of neurodegeneration, we targeted the *pyruvate kinase (pyk)* gene, a key enzyme in glycolysis that catalyzes the final step of glucose metabolism—converting phosphoenolpyruvate (PEP) to pyruvate while generating ATP. PyK is critical in sustaining metabolic homeostasis in neurons by ensuring a continuous supply of pyruvate to mitochondria for oxidative phosphorylation, thereby coupling glycolysis to oxidative phosphorylation and maximizing ATP yield. To this end, disrupting *pyk* expression in worms, flies, and mammalian neurons shifts energy metabolism towards glycolysis and decreases mitochondria bioenergetics, recapitulating metabolic imbalances observed in neurodegenerative diseases [35–37]; further, mice lacking pyruvate kinase postnatally in hippocampal neurons exhibit age-dependent learning and memory deficits [37].

In this study, we observed that loss of PyK induces progressive synaptic and axonal degeneration in motor neurons, carried out by the canonical neurodegenerative regulators Wnd/DLK and SARM1. We also found evidence that Wnd/DLK signaling activates a genetic program preceding synapse loss, via the activator protein 1 (AP-1) transcription factor Fos. When we challenged PyK-deficient axons with an additional stressor (injury) prior to synapse loss, we discovered that injury-induced axon degeneration is delayed, suggesting the presence of a neuroprotective mechanism. This delay shows dependence on Wnd/DLK and Fos, indicating that they genetically regulate a protective, axon integrity-conserving mechanism in PyK-deficient neurons. This delay in degeneration is also restored by overexpression of SARM1, the axonal localization of which we found to be significantly reduced in PyK-deficient neurons. This demonstrates that both the progressive synapse loss and delayed axon degeneration in PyK-deficient neurons are sensitive to SARM1 manipulation. This supports a rheostat model of Wnd/DLK, either promoting or inhibiting neurodegeneration (potentially through the regulation of SARM1) to control synapse survival or destruction during metabolic stress. Together, these findings provide new insights into how neurons rely on central carbon metabolism, particularly the coupling of glycolysis with oxidative phosphorylation via PyK, to support their projections and reveal a dynamic signaling response that determines synapse destruction or survival. Understanding these mechanisms provides a framework for advancing metabolic interventions as potential therapeutic strategies in neurodegenerative diseases and identifying early biomarkers of neuronal vulnerability and disease progression.

## Materials and Methods

*Animal Lines*. W118, UAS-*luciferase*-RNAi (RRID:BDSC_31603), UAS-*lexA*-RNAi (RRID:BDSC_67947), UAS-Luciferase (RRID:BDSC_35788), QUAS-gRNA (RRID:BDSC_67539), UAS-*pyk*-RNAi (RRID:BDSC_35218), UAS-*wnd*-RNAi (VDRC 103410), UAS-*dsarm*-RNAi (VDRC 105369), UAS-*akr1B*-RNAi (RRID:BDSC_67838), UAS-Fos^DN^ (RRID:BDSC_7214, [38]), UAS-dSarm-GFP (gift from Marc Freeman lab, [22]), D42-Gal4 [39], BG380-Gal4 [40], M12-Gal4 [41], *puc*-LacZ [42], PyK-sgRNA (RRID:BDSC_78770), UAS-SEpHluorin-FusionRed-Ras (UAS-pHusion-Ras, gift from Gregory Macleod lab, membrane-bound variant of pHusion [43]), UAS-Dcr2 (RRID:BDSC_24650), UAS-Cas9 (RRID:BDSC_58985).

### Animal Rearing

Flies were grown on yeast glucose food (10% baker’s yeast, 10% glucose, 0.3% tegosept, 0.44% proprionic acid, and 1.5% agar). Flies were maintained at 25°C in a humidity-controled incubator under a 12:12 hour light/dark cycle. 2^nd^ or 3^rd^ instar larvae were used for all experiments, as specified.

### Axon Degeneration Scoring

Axons were visualized using expression of fluorophore-tagged mCD8 and were scored using the following scale: 0% - completely continuous, 33% - continuous with varicosities, 66% - partially continuous and partially fragmented, 100% - fully fragmented, as previously described [31]. All scoring was done blinded.

### Synaptic Degeneration Scoring

Degeneration of NMJs was scored as the total number of breaks in the synaptic membrane visualized with anti-HRP antibodies or with expressed fluorophore-tagged mCD8, consistent with similar methods [31]. NMJs from segments 3-5 were scored, with up to six NMJs per animal. All scoring was done blinded.

### Larval Crawling Assay

Larvae were placed in a 1μL drop of 10% FD&C Blue #1 dye and given 60 seconds to move around a 10cm petri dish with a 2mm spaced grid. The larvae were then removed and the number of spaces positive for dye were counted. All assays were done blinded.

### Fluorescent Protein Quantifications

Neuron cell body quantifications were done by outlining neurons corresponding to segments 5-7 (six cells (three pairs) per larva using the M12-Gal4 driver) and summing the total signal in the selected channels (one total value per larva). The ROIs were then dragged to empty space beside the nerve cord for background readings. For axon quantifications, 20 μm diameter circular stamps were taken from the axons corresponding to the measured neuron cells bodies and summed for the same channels. Five stamps were taken from each pair of axons (for 15 stamps total per larva, averaged into one value per larva) and nine stamps of the same size were taken from empty spaces by the axons and averaged for a background reading. Plotted values are relative to the first control.

### JNK Signaling Reporter (puc-LacZ)

To test for activation of kinase pathways involving JNK, the *puc*-LacZ report [44] was used with the pan-motor neuron driver BG380-Gal4. Male progeny were used as the Gal4 driver is on the X chromosome. Motor neurons along the midline of the nerve cord corresponding to segments 4-7 of the larvae (8-10 cells per segment, 32-40 per animal) were quantified and averaged for one value per animal. Background measurements were taken to the side of each group of neurons (3 per segment, 12 per animal) and averaged. Plotted values are relative to the first control.

### Nerve Injury

Larvae were placed in ice-cold phosphate-buffered saline (PBS) for 15 minutes before injury. Larvae were then transferred to an upside-down petri dish and were pinched with Dumostar number 5 forceps on the ventral side (just posterior of the nerve cord), crushing the nerves and severing the neurites within. Larva were then placed on small petri dishes with fly food to recover. A more detailed protocol with images is available [31].

### Dissections, Tissue Preparation, and Antibody Staining

Larvae were dissected in an ice bath in PBS. Dissected tissue was fixed in room temperature (RT) 4% paraformaldehyde in PBS for 20 minutes, then washed three times quickly with PBS. Tissue samples were then blocked for 1 hour at RT with 5% normal goat serum (NGS) in PBS with 0.25% Triton X detergent (PBST). Primary antibodies (when used) were diluted in 5% NGS in PBST at concentrations of 1:1000 (Rat anti-mCD8 [Invitrogen MCD0800]) or 1:100 (Mouse anti-CSP [DSHB AB49] and Mouse anti-β-Galactosidase [DSHB 40-1A]) and incubated with fixed tissue overnight at 4C on a low shaker setting (~30 RPM). If primary antibodies were used, tissue samples were washed with PBST three times for 10 minutes each at RT before moving to secondary staining. Secondary (and conjugated) antibodies were diluted 1:1000 (488 Goat anti-Mouse (Fisher), 488 Rabbit ant-GFP (Fisher), Cy3 Goat anti-HRP (Fisher), Cy3 Goat anti-Rat (Fisher)) in 5% NGS in PBST and were incubated with tissue samples for 2 hours at RT on a low shaker setting (~30 RPM). Tissue samples were then washed three times for 10 minutes with PBST before being mounted on glass slides using Prolong Diamond mounting media. Samples were given at least 24 hours to set before being imaged.

### Microscopy

Images were taken on an Improvision spinning disk confocal and a Stellaris Leica microscope. All conditions in a given repeat for experiments comparing fluorescence levels were captured in the same imaging session and using the same laser and exposure settings. Quantifications and image processing were done using the Volocity and ImageJ softwares.

### Dominant-Negative Fos

To suppress Fos activity in several experiments, the Fos^DN^ construct was used [38], which contains the DNA binding domain and ability to dimerize with other transcription factors [45], but missing the section required for activation of transcription [46].

### Gene Ontology Term Generation

Published RNA sequencing data from [36] were analyzed with the g:Profiler tool [47] with inclusion criteria being all upregulated genes with an adjusted p-value (padj) of less than 0.05.

## Results

### Loss of pyruvate kinase (PyK) leads to progressive synaptic and axonal degeneration in motor neurons

Metabolic homeostasis perturbations have been widely observed in both peripheral and central nervous system disorders, suggesting that tight metabolic regulation is crucial for neuronal health [6–10]. However, whether these metabolic changes are a cause or consequence of neurodegeneration remains unresolved. To directly test the effects of metabolic disruption on neuronal integrity, we examined how impairing metabolic homeostasis influences motor neuron projections in vivo.

We hypothesized that disrupting metabolic homeostasis would lead to axonal loss and synaptic degeneration. To induce a controlled metabolic perturbation, we used specific loss-of-function and partial loss-of-function genetic manipulations in motor neurons of fruit fly larvae. We selected the PyK gene as our target because as the last rate-limiting enzyme of glycolysis, it sits at a critical metabolic nexus, coupling glycolysis to the tricarboxylic acid (TCA) cycle and oxidative phosphorylation ([48,49] and diagram in **Figure 1A**). Disruptions in pyruvate kinase activity have been shown to alter metabolic homeostasis across multiple species, from invertebrates to mammals [35,36,49–51]. The fly PyK gene shares >60% sequence identity with human pyruvate kinase (PKM), and its loss in flies leads to glycolytic disruption and a reduction in TCA cycle activity [36]. Specifically, fly *pyk* mutants phenocopy other glycolytic mutants, exhibiting a severe block in glycolysis, and accumulate intermediates upstream of pyruvate, such as 2-phosphoglycerate, 3-phosphoglycerate and phosphoenolpyruvate, and depletion of fumarate, malate, lactate, 2-hydroxyglutarate (2HG), and proline [36]. Beyond the shared metabolic disruptions observed across species, PyK/PKM loss in both flies and mammals leads to cognitive impairments, including deficits in learning and memory [52–55]. Thus, manipulating PyK function in flies provides a direct and evolutionarily relevant model to study metabolic homeostasis disruption in the nervous system, with clear parallels to mammalian systems.

**Figure 1:**
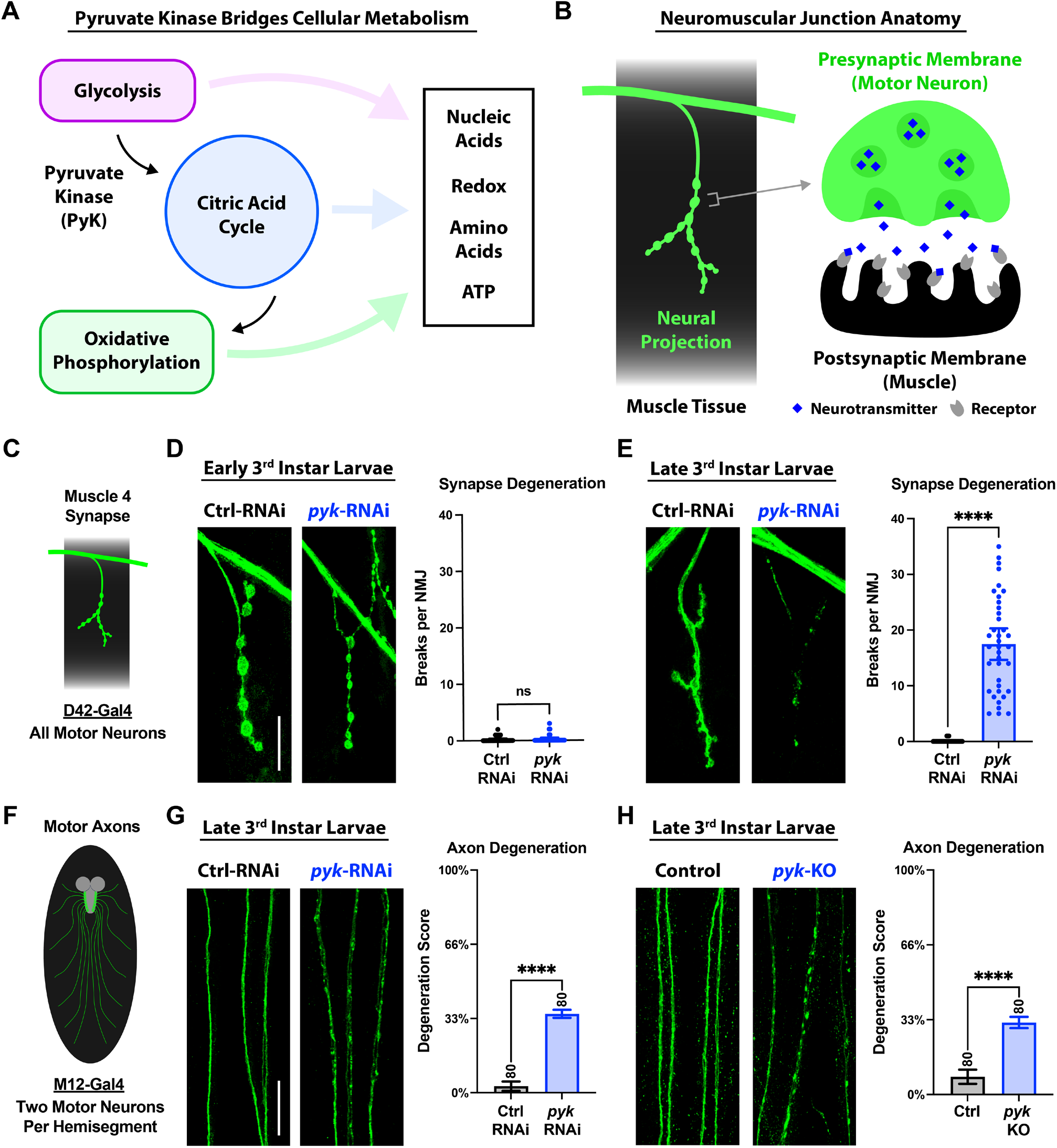
Loss of pyruvate kinase (PyK) causes progressive synaptic and axonal degeneration in motor neurons. **A)** Position of pyruvate kinase in the cellular metabolic hub of glycolysis, the citric acid cyclic, and oxidative phosphorylation. **B)** Schematic of the neuromuscular junction, which is made up of a motor neuron axon projecting onto the muscle membrane and forming a synapse. Release of neurotransmitters at this synapse triggers muscle contraction. **D)** All synapses imaged in this study are at Muscle 4 of the larval body wall, using the pan-motor neuron driver D42-Gal4 for genetic manipulations. **E)** NMJs of early 3^rd^ instar larvae expressing *luciferase*-RNAi or *pyk*-RNAi in motor neurons, with co-expression of the mCD8-GFP membrane marker for visualization. Between 2-6 Muscle 4 NMJs per animal (innervating body segments 3-5) from 8 larvae per condition were scored blinded for number of breaks in the synaptic membrane and plotted individually. **F)** NMJs of late 3^rd^ instar larvae expressing *luciferase*-RNAi or *pyk*-RNAi in motor neurons, with mCD8-GFP for visualization. Between 3-6 Muscle 4 NMJs per animal (innervating body segments 3-5) from 8 larvae per condition were scored blinded for number of breaks in the synaptic membrane and plotted individually. **G**) Axons shown in this study are of those labeled by M12-Gal4, which expresses in two motor neurons per larval hemisegment. **H**) Motor neuron axon pairs of late 3^rd^ instar larvae expressing *luciferase*-RNAi or *pyk*-RNAi, with mCD8-GFP for visualization. 10 axon pairs (innervating body segments 3-7) each from 8 larvae per condition were scored blinded for extent of degeneration using a 4-point scale (0% - completely continuous, 33% - continuous with varicosities, 66% - partially continuous and partially fragmented, 100% - fully fragmented). **I**) Motor neuron axon pairs of control (wildtype) or *pyk*-gRNA late 3^rd^ instar larvae expressing Cas9 for guided knockout with mCD8-RFP for visualization (colored green). 10 axon pairs (innervating body segments 3-7) each from 8 larvae per condition were scored blinded for extent of degeneration using a 4-point scale (0% - completely continuous, 33% - continuous with varicosities, 66% - partially continuous and partially fragmented, 100% - fully fragmented). All statistical comparisons are unpaired t-tests. Scale bars are 20 μm. Significance symbols: * p<0.05, ** p<0.01, *** p<0.001, **** p<0.0001, ns = not significant.

To test the hypothesis that PyK loss-induced metabolic disruptions contribute to alterations in axonal and synaptic integrity, we knocked down this gene in the fly motor neurons using the Gal4/UAS system, which targets genetic manipulations with spatial specificity. We evaluated synaptic integrity at the neuromuscular junction (NMJ), where the motor neuron axon projects out from the nerve to form a synapse onto the muscle membrane (diagram in **Figure 1B**). We first compared the effects of pan-motor neuronal *pyk* knockdown (KD) on synaptic integrity (specifically at the well-characterized larval muscle 4) using the established D42-Gal4 motor neuron driver and co-expressing the membrane-bound mCD8-GFP protein ([31,39,56,57] and diagram in **Figure 1C**). Control flies expressing a luciferase double-stranded RNA (dsRNA) transgene for RNA interference (RNAi) in the motor neurons showed robust synaptic structures in both early and late 3rd instar larval stages (about 2 days apart) (**Figure 1D-E**). However, *pyk* RNAi resulted in synaptic degeneration by late 3rd instar, as shown by the many breaks observed across the synaptic membrane (**Figure 1D-E**). This phenotype is similar in strength to the full synaptic fragmentation reached by larval NMJs following nerve injury [31]. Thus, *pyk* knockdown motor neurons are capable of initially forming synapses (as viewed at early 3rd instar), which later degenerate (by late 3rd instar). These early synapses, however, have altered morphology as revealed through the synaptic cysteine string protein (CSP), a marker for synaptic vesicles, which was reduced in *pyk* knockdown compared to control animals (**Supplemental Figure 1**). Consistent with the structural data, these animals show strong impairments in locomotion when tested on a motility behavioral assay (**Supplemental Figure 2**). We next examined the integrity of the motor neuron axons using the more selective driver M12-Gal4, which expresses in two motor neurons per larval hemisegment, allowing for genetic manipulations that may not be tolerable when done in the full motor nervous system ([31,41] and diagram in **Figure 1F**). This allowed us to clearly score individual axonal structures within nerves and determine a total degeneration score based on the amount of varicosities and fragmentation of these neurites (see methods). Compared to control axons, *pyk* knockdown axons were continuous, but marked with varicosities throughout, consistent with early stages of axon degeneration (**Figure 1G**). This phenotype was confirmed via short guide RNA (sgRNA)-guided Cas9-mediated knockout of *pyk*, which also results in pronounced axonal varicosities (**Figure 1H**). Together these results indicate that perturbing metabolic homeostasis in the motor neurons leads to progressive synaptic and axonal degeneration and locomotor defects.

**Figure 2:**
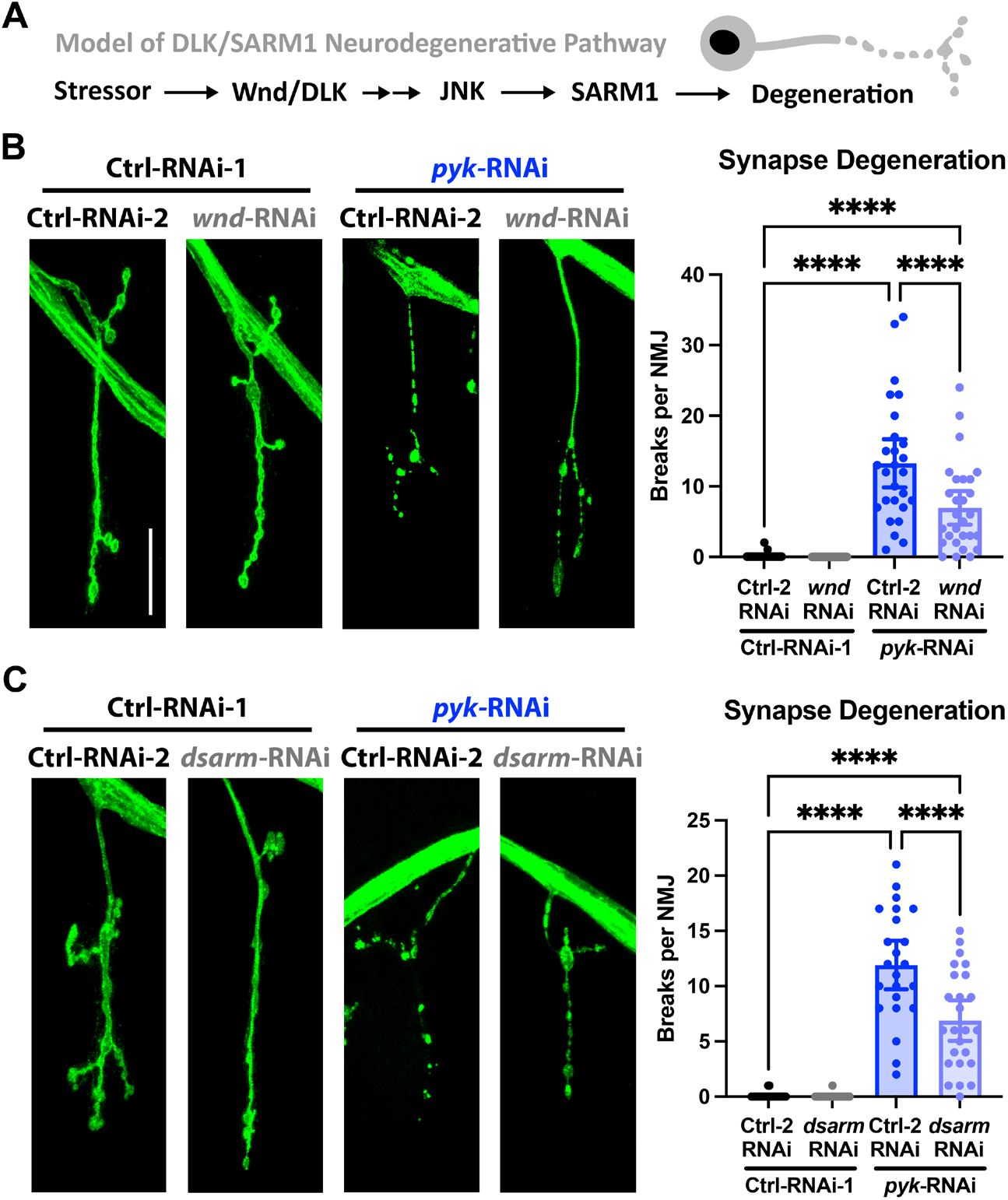
Synaptic degeneration due to PyK deficiency requires Wnd/DLK and dSarm/SARM1. **A)** Schematic of Wnd/DLK signaling and SARM1 regulation in axon degeneration. **B)** NMJs of late 3^rd^ instar larvae expressing *luciferase*-RNAi or *pyk*-RNAi with *lexA*-RNAi or *wnd*-RNAi in motor neurons (D42-Gal4), with Cy3 anti-HRP staining for visualization (neuronal membrane, colored green), scored blinded for breaks in the synaptic membrane. Between 3-6 Muscle 4 NMJs per animal (innervating body segments 3-5) from 6 larvae per condition were scored blinded for number of breaks in the synaptic membrane and plotted individually. **C)** NMJs of late 3^rd^ instar larvae expressing *luciferase*-RNAi or *pyk*-RNAi in combination with *lexA*-RNAi or *wnd*-RNAi in motor neurons (D42-Gal4), with Cy3 anti-HRP staining for visualization (neuronal membrane, colored green). Between 1-5 Muscle 4 NMJs per animal (innervating body segments 3-5) from 7-8 larvae per condition were scored blinded for number of breaks in the synaptic membrane and plotted individually. All statistical comparisons are one-way ANOVA with Tukey correction for multiple comparisons. Scale bars are 20 μm. Significance symbols: * p<0.05, ** p<0.01, *** p<0.001, **** p<0.0001, ns = not significant.

### Synaptic degeneration due to PyK deficiency requires the canonical Wnd/DLK and dSarm/SARM1 pathways

Having established that manipulations of PyK are sufficient to induce synaptic and axonal degeneration, we next set to identify effectors of this neurodegeneration. We focused on canonical degenerative signaling pathways to understand the potential cross-talk between metabolic alterations and disease-relevant signaling. The MAP3K DLK (Wnd in flies) pathway is a well-known and evolutionarily-conserved axon degeneration, regeneration, and protection effector in several disease models and injury paradigms [12,33,58–61]. Further, SARM1 (dSarm in flies) has risen to the forefront of neurodegeneration research following the discovery of its local role in promoting axon degeneration through the breakdown of the electron carrier nicotinamide adenine dinucleotide (NAD) [22,23]. This activity is allosterically regulated by the ratio of NAD and its precursor nicotinamide mononucleotide (NMN), which is maintained axonally by the neuroprotective nicotinamide mononucleotide adenylyl transferase (NMNAT), placing SARM1 as a sensor for the metabolic disruption linked to axonal and synaptic degeneration [20,62–68]. Importantly, Wnd/DLK and related MAPK signaling actors have been connected to the regulation of SARM1 in neurodegenerative contexts ([11–17,69,70], abbreviated schematic in **Figure 2A**). Intriguingly, fly *pyk* mutants show changes in the expression of genes involved in synaptic and axonal function and MAPK signaling, including those related to neurodegeneration such as Wnd/DLK and SARM1 and their downstream effectors, which are upregulated ([36] and **Supplemental Figure 3**).

**Figure 3:**
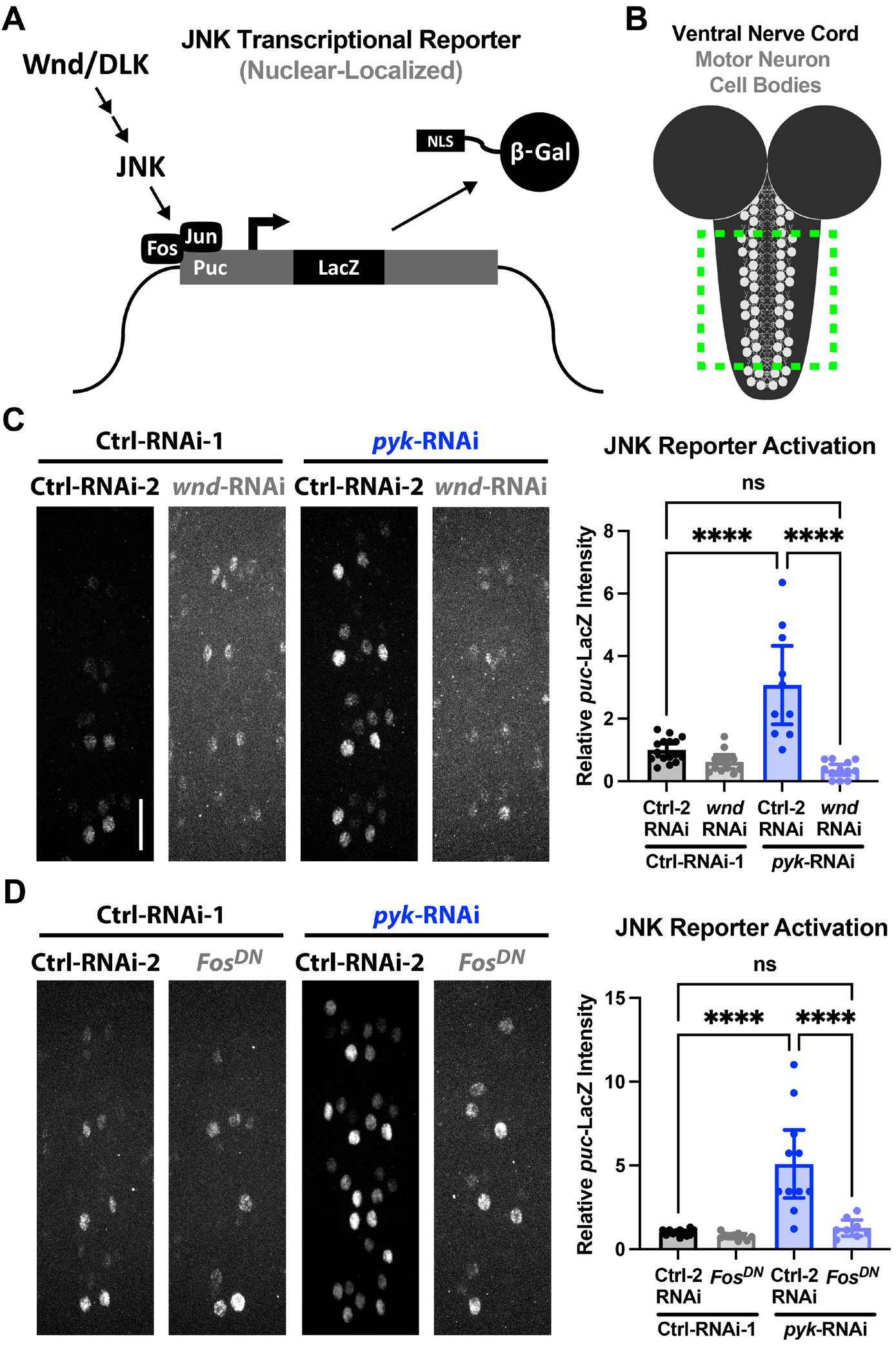
Knockdown of PyK activates Wnd/DLK signaling to the nucleus via Fos. **A)** Mechanism of the JNK genetic reporter used in this study (*puc*-LacZ, [42]), which produces β-Galactosidase in response to JNK-dependent activation of the Fos/Jun (AP-1) transcription factors. **B)** The *puc*-LacZ reporter output (β-Gal with nuclear localizing signal) is stained and measured at the midline motor neurons of the ventral nerve cord, using the pan-motor neuron driver BG380-Gal4 for genetic manipulations. **C)** Levels of β-Gal in the midline motor neuron nuclei of *puc*-LacZ mid 3^rd^ instar larvae expressing either *luciferase*-RNAi or *pyk*-RNAi in combination with either *lexA*-RNAi or *wnd*-RNAi. Nuclei from motor neurons innervating body segments 4-7 (8-10 cells per corresponding segment, 32-40 total per animal) from 10-16 larvae per condition were quantified for β-Gal staining intensity and were plotted as animal averages. **D)** Levels of β-Gal in the midline motor neuron nuclei of *puc*-LacZ mid 3^rd^ instar larvae expressing either *luciferase*-RNAi or *pyk*-RNAi in combination with either *lexA*-RNAi or Fos^DN^. Nuclei from motor neurons innervating body segments 4-7 (8-10 cells per corresponding segment, 32-40 total per animal) from 8-12 larvae per condition were quantified for β-Gal staining intensity and were plotted as animal averages. All statistical comparisons are one-way ANOVA with Tukey correction for multiple comparisons. Scale bars are 20 μm. Significance symbols: * p<0.05, ** p<0.01, *** p<0.001, **** p<0.0001, ns = not significant.

To answer these questions, we knocked down Wnd/DLK or dSarm/SARM1 in control or PyK-deficient motor neurons and examined the level of synaptic degeneration. We reasoned that if these critical effectors were involved, then there should be a decrease in the severity of the synaptic phenotypes. In PyK-deficient motor neurons, knockdown of either Wnd/DLK or dSarm/SARM1 led to a ~50% suppression of synaptic fragmentation (**Figure 2B-C**). These data indicate that Wnd/DLK and SARM1 are necessary to mediate the synaptic degeneration observed with *pyk*-induced dysfunctions in motor neurons. They also indicate that these pathways are genetically downstream of the metabolic disruptions driven by PyK deficiencies.

### PyK deficiency activates a transcriptional response dependent on AP-1 and Wnd/DLK

Upon activation, Wnd/DLK initiates a kinase signaling cascade that engages JNK and the downstream AP-1 transcription factors Fos (Kay in flies) and Jun (Jra in flies) to activate genetic responses involved in axon regeneration, axon degeneration, synaptic growth, and apoptosis, depending on context [33,60,71–76]. AP-1 is also implicated indirectly and directly in the regulation of metabolic genes [77–79]. Because the axonal and synaptic degeneration of PyK-deficient neurons requires Wnd/DLK signaling, we sought to determine whether this pathway engages canonical Wnd/DLK transcriptional responses and effectors. To address this, we used the genetically encoded, nuclear-localized puckered (*puc*)-LacZ reporter, which allows visualization of Wnd/DLK-induced transcription in motor neuron nuclei ([44] and diagrams in **Figure 3A-B**). Puc encodes a dual-specificity phosphatase (DUSP) that negatively regulates JNK signaling, providing a well-characterized biosensor of JNK activity [42]. The expression of this gene is widely used as a readout of JNK activation and AP-1-dependent transcription [42,59,80–83]. In PyK-deficient motor neurons, *puc*-LacZ expression was significantly increased, indicating robust activation of the canonical Wnd/DLK-JNK-AP-1 signaling cascade (**Figure 3C-D**). However, this effect was entirely dependent on Wnd/DLK and Fos, as Wnd/DLK knockdown or expression of a dominant-negative Fos allele abolished the *pyk*-induced increase in *puc*-LacZ expression (**Figure 3C-D**). These findings demonstrate that PyK deficiency activates a transcriptional response downstream of DLK, which requires the Fos transcription factor. Consistent with these genetic observations, reanalysis of transcriptomic data from Heidarian et al. 2023 revealed that genes altered by *pyk* mutations were significantly enriched for the AP-1 binding motif (TF:M00199_1, NTGASTCAG, padj=2.03E-03), a sequence recognized by the Fos/Jun complex ([36] and **Supplemental Figure 3**). Among these genes, *puc* mRNA exhibited a strong upregulation (log2 fold change=0.444, padj=6.00E-04), further supporting the conclusion that PyK loss triggers a transcriptional program driven by JNK signaling through the canonical AP-1 factors [36].

### Neuroprotective responses to PyK deficiency revealed by Wallerian Degeneration

Although the coupling between glycolysis and the TCA cycle is essential for neuronal health, recent studies suggest this relationship may be context-dependent and involve non-autonomous mechanisms [84–89]. In healthy neurons, metabolic homeostasis sustains the high energetic demands of synaptic transmission, ion homeostasis, and overall neuronal stability [1,9,90,91]. Upregulation of glycolysis can be protective in models of ALS, Alzheimer’s disease, and Parkinson’s disease [34,92,93]. In contrast, reduction of glycolytic flux can be beneficial in models of neuroinflammation and traumatic brain injury (TBI) [94,95]. In addition to the impact of glucose metabolism itself on neuron survival, neuroprotective stress responses could be activated by disruption of this pathway, which could be in opposition to the neurodegenerative signaling activated by PyK deficiency. Evidence for this possibility can be seen with axon injury, where an initial injury (conditioning lesion) can activate a protective stress response that inhibits degeneration following a second injury, with Wnd/DLK signaling and the downstream transcription factor Fos capable of establishing this protection ([58,96,97] and diagrams in **Figure 4A-B**). This is evidence that genetic responses to neuronal stressors can boost axon integrity, which can then be revealed and measured through axon injury. Based on this knowledge, we tested the hypothesis that *pyk* knockdown may confer neuroprotection that could be revealed through axon injury, potentially mediated by Wnd/DLK-JNK-AP-1 signaling.

**Figure 4:**
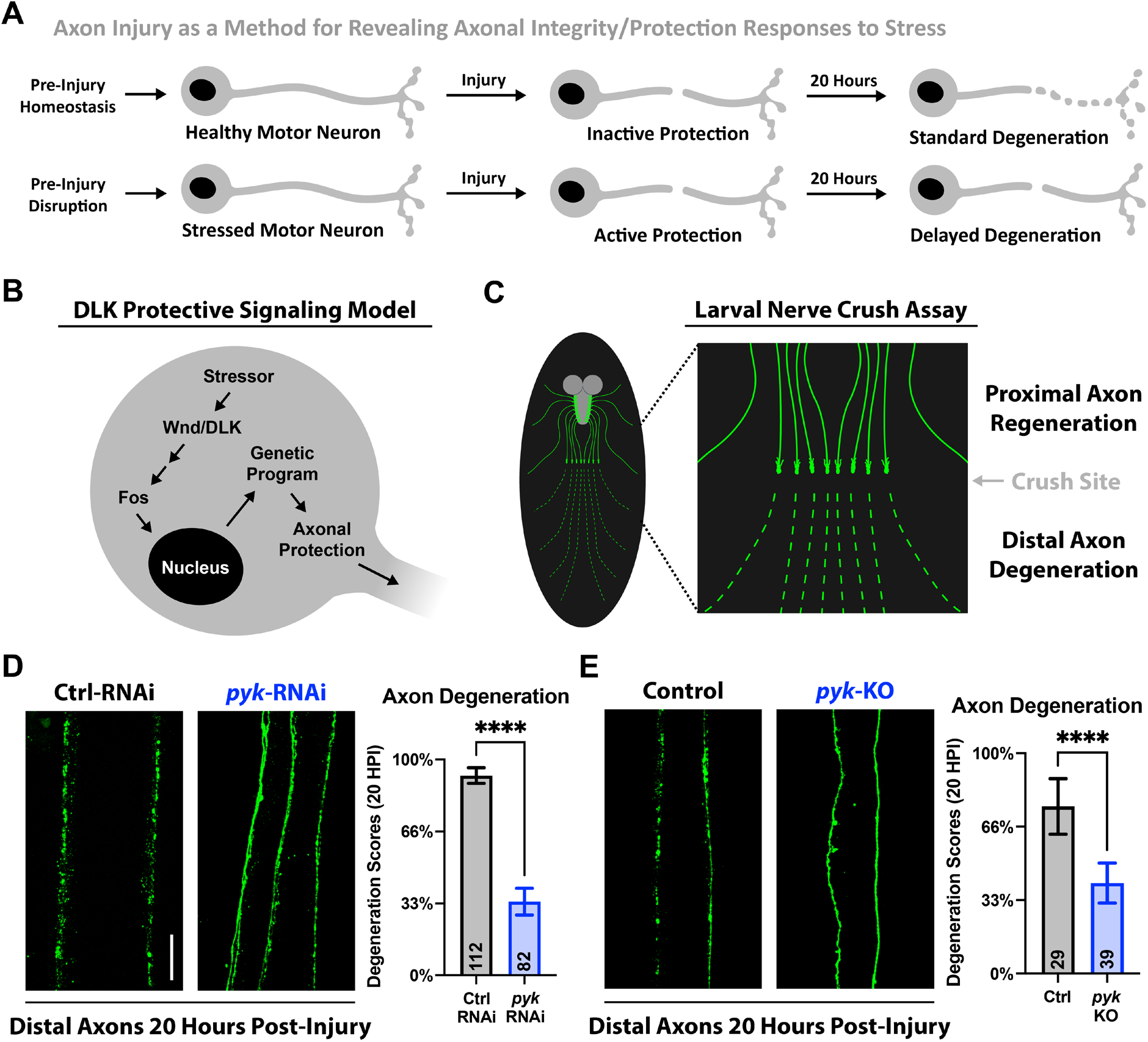
Loss of PyK delays injured axon degeneration (Wallerian degeneration). **A)** Model of axon protection based on pre-injury stress response (such as in a conditioning lesion [58]). **B)** Wnd/DLK neuroprotective signaling diagram, content reviewed in [33]. **C)** Schematic of larval nerve crush and injury response, full protocol in [31]. **D)** Distal injured motor neuron axon pairs of mid 3^rd^ instar larvae expressing *luciferase*-RNAi or *pyk*-RNAi, with mCD8-GFP for visualization. Between 4-9 severed axon pairs each from 11-16 larvae per condition were scored blinded for extent of degeneration using a 4-point scale (0% - completely continuous, 33% - continuous with varicosities, 66% - partially continuous and partially fragmented, 100% - fully fragmented). Total number of axons pairs scored per condition shown on graph. **E)** Distal injured motor neuron axon pairs of control (*QUAS*-gRNA) or *pyk*-gRNA mid 3^rd^ instar larvae expressing Cas9 for guided knockout, with mCD8-GFP for visualization. Between 5-8 severed axon pairs each from 4-6 larvae per condition were scored blinded for extent of degeneration using a 4-point scale (0% - completely continuous, 33% - continuous with varicosities, 66% - partially continuous and partially fragmented, 100% - fully fragmented). Total number of axons pairs scored per condition shown on graph. All statistical comparisons are unpaired t-tests. Scale bar is 20 μm. Significance symbols: * p<0.05, ** p<0.01, *** p<0.001, **** p<0.0001.

To investigate the impact of *pyk* knockdown on injury-induced Wallerian axon degeneration, we used a well-established larval nerve crush model to measure motor axon fragmentation 20 hours post-injury ([31] and diagram in **Figure 4C**). While the severed axons of early 3rd instar control animals exhibited rapid and complete degeneration, *pyk*-RNAi knockdown or gRNA-directed Cas9-mediated *pyk* knockout significantly delayed axonal self-destruction (**Figure 4D-E**). Thus, disrupting glycolytic flux through PyK genetic manipulation delays the breakdown of axons severed from the cell body.

Given the role of Wnd/DLK-JNK-AP-1 in the axon protection following a conditioning lesion, we next examined whether Wnd/DLK and its nuclear signaling mediated the protective effects of *pyk* knockdown ([58,96] and diagram in **Figure 5A**). Extensive work examining this pathway has revealed its multiple, context-dependent roles in axon growth, degeneration, regeneration, and protection [12,15,32,33,58–61,74,98].

**Figure 5:**
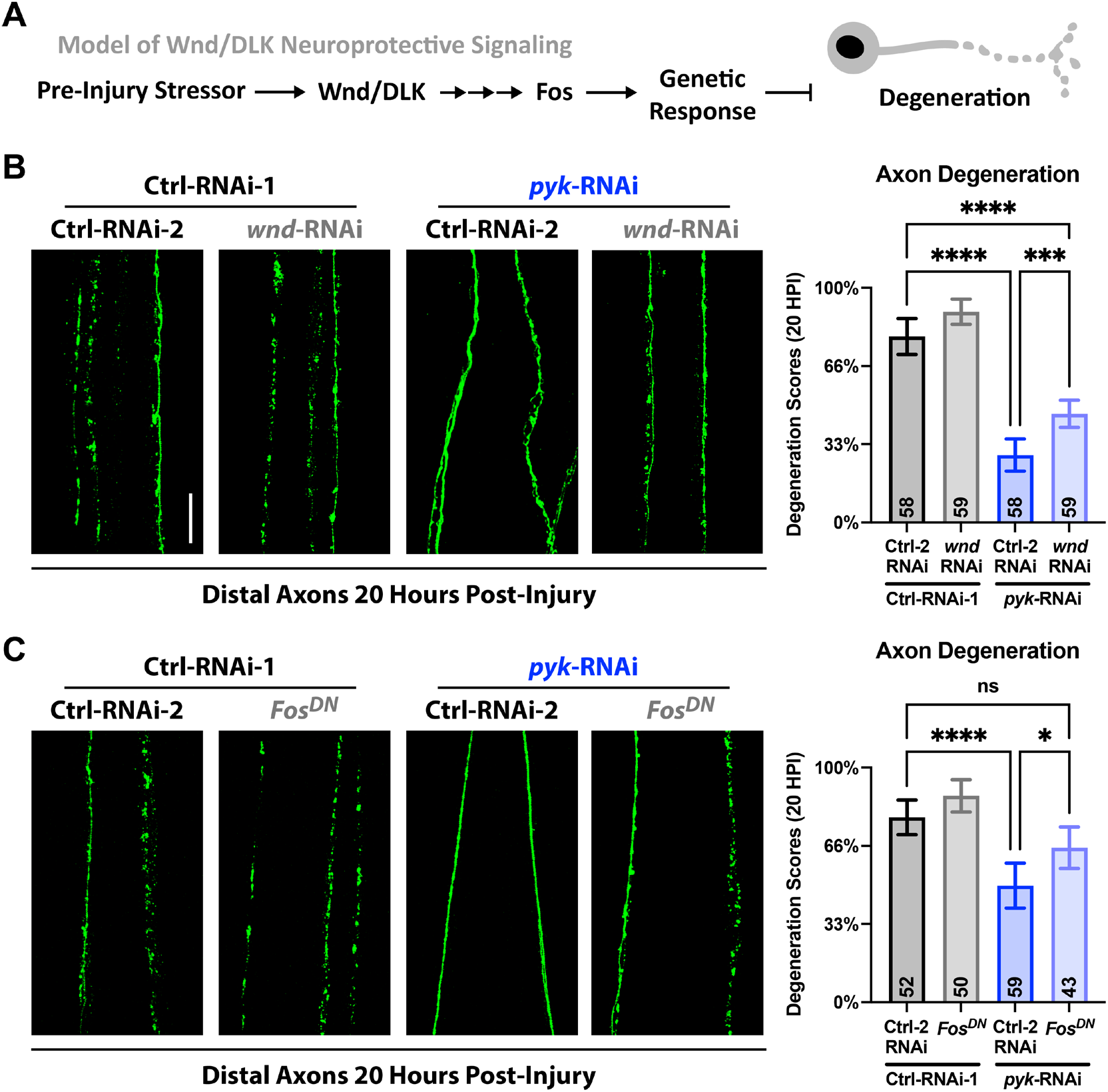
Delayed Wallerian degeneration of PyK knockdown shows dependence on Wnd/DLK signaling. **A)** Proposed model being tested of Wnd/DLK neuroprotective signaling inducing a genetic response that inhibits SARM1 to delay axon degeneration. **B)** Distal injured motor neuron axon pairs of mid 3^rd^ instar larvae expressing *luciferase*-RNAi or *pyk*-RNAi in combination with either *lexA*-RNAi or *wnd*-RNAi, with mCD8-GFP for visualization. Between 5-10 severed axon pairs each from 8 larvae per condition were scored blinded for extent of degeneration using a 4-point scale (0% - completely continuous, 33% - continuous with varicosities, 66% - partially continuous and partially fragmented, 100% - fully fragmented). Total number of axons pairs scored per condition shown on graph. **C)** Distal injured motor neuron axon pairs of mid 3^rd^ instar larvae expressing *luciferase*-RNAi or *pyk*-RNAi in combination with either *lexA*-RNAi or Fos^DN^, with mCD8-GFP for visualization. Between 2-8 severed axon pairs each from 8 larvae per condition were scored blinded for extent of degeneration using a 4-point scale (0% - completely continuous, 33% - continuous with varicosities, 66% - partially continuous and partially fragmented, 100% - fully fragmented). Total number of axons pairs scored per condition shown on graph. All statistical comparisons are one-way ANOVA with Tukey correction for multiple comparisons. Scale bar is 20 μm. Significance symbols: * p<0.05, ** p<0.01, *** p<0.001, **** p<0.0001, ns = not significant.

Following damage to neurons, Wnd/DLK nuclear signaling pathways promote both protection and regeneration through transcriptional changes [33,58,71– 74,96]. Within the distal stump, the Wnd/DLK downstream target JNK promotes Wallerian degeneration, an active, local degenerative processes that drives the destruction of the severed axon [99,100]. The role of transcriptional regulation in the priming of axons for post-injury degeneration, including by Wnd/DLK, is still being examined [33,99–101]. We found that co-knockdown of Wnd/DLK in PyK-deficient motor neurons partially restored neurodegeneration, albeit not at the same levels of control animals (**Figure 5B**). Inhibition of the nuclear arm of Wnd/DLK signaling through the expression of dominant-negative Fos also rescued the delayed degeneration (**Figure 5C**). These genetic interactions suggest that the protective responses to perturbed glycolysis require, at least in part, genetic regulation by the Wnd/DLK-JNK-AP-1 pathway. However, it is important to note that the rescue of neurodegeneration speed was incomplete. This could be a technical limitation of using partial knockdown and dominant negative alleles of Wnd/DLK and Fos, or could implicate the involvement of additional pathways. We tested one of these potential pathways through co-knockdown of PyK with adenosine monophosphate activated protein kinase (AMPK), a well-characterized regulator of cellular energy that is activated by disruption of glycolysis and has been linked to neuroprotective mechanisms [102–106], but found no significant impact on either *puc*-LacZ JNK reporter expression or delayed axon degeneration post-injury (**Supplemental Figure 4**).

As the contributions of Wnd/DLK and Fos to the delayed Wallerian degeneration of PyK-deficient axons are expected to be through genetic regulation, and given the partial rescue of degeneration speed when Wnd/DLK is targeted, we next searched for local changes within the axons themselves that could reveal the mechanism of this protection. Following axon injury, local degeneration of the distal stump requires SARM1, whose localization to the axon has been implicated in delayed Wallerian degeneration ([17,22,23] and diagram in **Figure 6A**). Using overexpression of GFP-tagged dSarm/SARM1, we found that dSarm/SARM1 levels are reduced specifically in the axons of PyK-deficient motor neurons compared to controls (**Figure 6B-C**). Based on this information, we would expect that overexpression of this effector could restore axon degeneration post-injury in PyK-deficient axons. Indeed, the delayed Wallerian degeneration seen in *pyk* knockdown was entirely abolished by dSarm/SARM1 overexpression (**Figure 6D**). Thus, the resilience to Wallerian degeneration in PyK-deficient neurons may involve a limitation on transport of SARM1 to the axon or increased turnover in this cellular extension.

**Figure 6:**
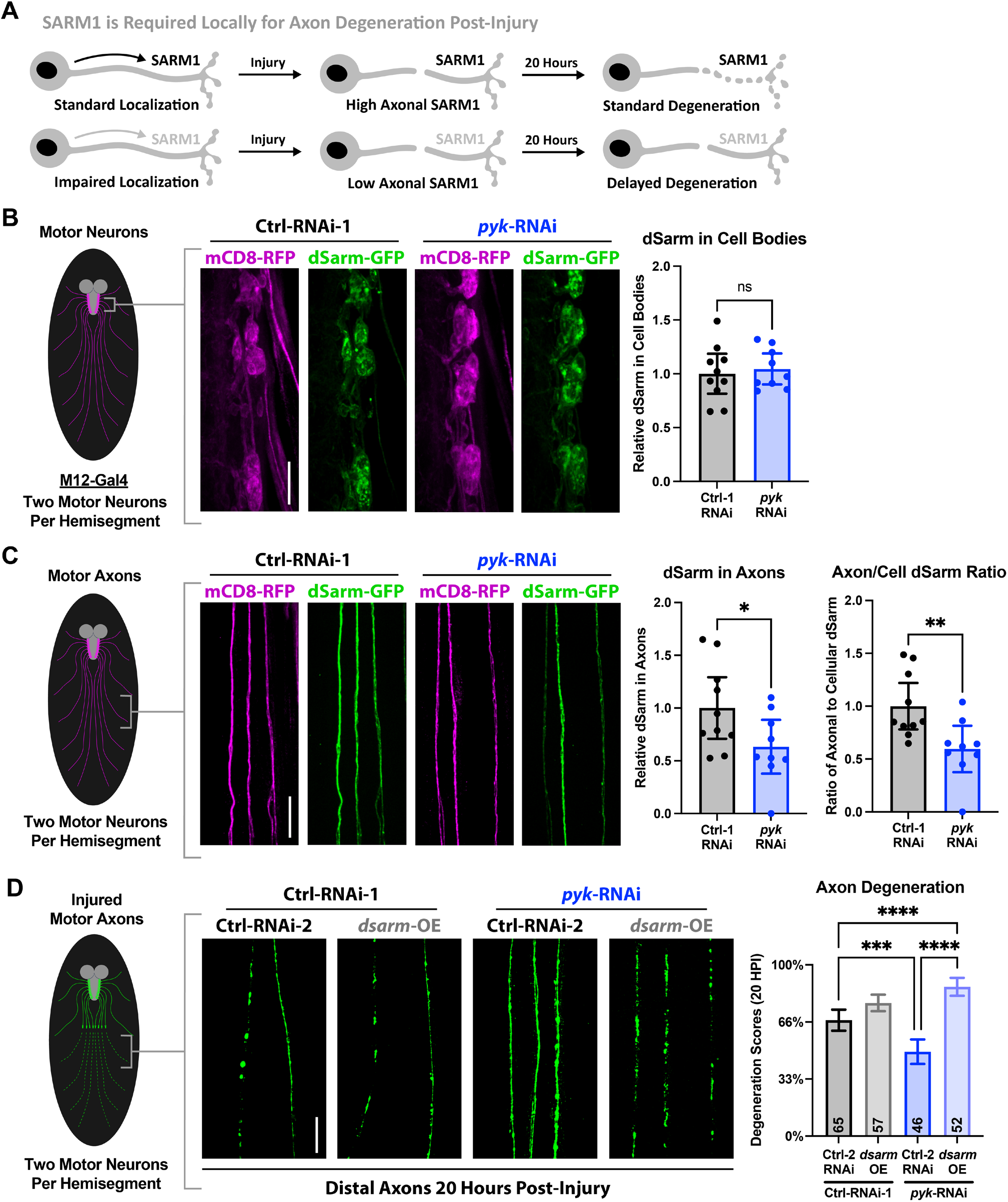
dSarm/SARM1 is decreased in PyK-deficient motor axons and fully restores degeneration when overexpressed. **A)** Model of the impact of SARM1 axonal localization on post-injury degeneration (supported by previous work [17,22,23]). **B)** Motor neuron cell bodies in early 3^rd^ instar larvae expressing *luciferase*-RNAi or *pyk*-RNAi with co-expressed GFP-tagged dSarm/SARM1 driven by M12-Gal4. The total signal of GFP-dSarm (green) per 3 pairs of M12-Gal4-expressing neurons corresponding to larval hemisegments 5-7 (6 neurons total per animal) was summed and plotted relative to the sum of the co-expressed membrane marker mCD8-RFP (magenta) for 9-10 larvae per condition. Data points represent animal averages. **C)** Axons corresponding to motor neurons from (A). The level of GFP-dSarm relative to mCD8-RFP was calculated and animal averages were plotted alone (left graph) or relative to the neuron cell body ratio (right graph) to calculate axon localization for 9-10 larvae per condition. **D)** Distal motor neuron axon pairs of mid 3^rd^ instar larvae expressing *luciferase*-RNAi or *pyk*-RNAi in combination with either *lexA*-RNAi or dSarm, with mCD8-RFP (colored green) for visualization. Between 2-8 severed axon pairs each from 8-9 larvae per condition were scored blinded for extent of degeneration using a 4-point scale (0% - completely continuous, 33% - continuous with varicosities, 66% - partially continuous and partially fragmented, 100% - fully fragmented). Total number of axons pairs scored per condition shown on graph. Statistical comparisons in **B** and **C** are unpaired t-tests. Statistical comparisons in **D** are one-way ANOVA with Tukey correction for multiple comparisons. Scale bars are 20 μm. Significance symbols: * p<0.05, ** p<0.01, *** p<0.001, **** p<0.0001, ns = not significant.

## Discussion

Metabolic dysfunction has been widely observed across neurodegenerative diseases, yet the extent that these changes drive degeneration versus arise as a consequence of neuronal stress remains unresolved [6–10]. Glial dysfunction or axon demyelination can cause progressive degeneration in multiple organisms and disease models, implicating loss of external metabolic support in neurodegenerative mechanisms [6,87,107–111]. How neuron-specific perturbations of metabolism are linked to neurodegeneration and interact with conserved neurodegenerative and regenerative pathways has remained unclear. Here, by leveraging the well-characterized Drosophila NMJ model, we provide evidence that genetic perturbations in neuronal metabolic homeostasis directly activate neurodegenerative signaling pathways. Our findings demonstrate that targeting metabolic flux through glycolysis, the TCA cycle, and ultimately oxidative phosphorylation by PyK disruption impairs axonal and synaptic integrity, engaging the Wnd/DLK-JNK-AP-1 and dSarm/SARM1 pathways to drive degeneration. We also uncover a neuroprotective response to metabolic disruption, seen in delayed injury-induced Wallerian degeneration of PyK-deficient axons, an effect dependent on Wnd/DLK’s nuclear response and fully abolished by SARM1 overexpression. Together, these results suggest that metabolic state influences the balance between axon degeneration and survival through canonical neurodegenerative pathways, positioning glycolysis and its downstream metabolic effectors as key regulators of neuronal health.

### Metabolic perturbations directly engage neurodegenerative signaling via canonical pathways

Metabolism has emerged as central to understanding neurodegenerative diseases. Disorders from dementias to peripheral neuropathies have been linked to perturbations in metabolism that includes sugar, fat, and protein breakdown, and multiple anabolic pathways [107,112–121]. There remains difficulty in determining what aspects of these metabolic shifts may drive mechanisms of these conditions (versus existing as consequences), especially where metabolic pathways intersect. To better understand this, we genetically targeted metabolic homeostasis through depletion of PyK, an essential metabolic node that connects glycolysis with the citric acid cycle and oxidative phosphorylation. Following this metabolic perturbation in motor neurons, we found progressive synaptic and axonal degeneration. This degeneration requires both the MAP3K Wnd/DLK and the NADase SARM1, both of which are well-known neurodegenerative actors linked to a range of central and peripheral nervous system diseases, such as chemotherapy-induced peripheral neuropathy (CIPN), ALS, and Alzheimer’s disease [26,122–134]. SARM1, especially, has been tied to metabolic disruption during axon degeneration through its destruction of the electron carrier NAD [23]. As SARM1 is itself allosterically regulated by the balance of NAD and its precursor NMN [20,21], which may act as indicators of metabolic balance, there is reason to hypothesize SARM1 would be sensitive to metabolic disruptions separate from the nervous system injury response for which it is best known. By targeting metabolism genetically, our work suggests SARM1 and Wnd/DLK are capable of responding to metabolic perturbation in neurons. In addition to the implications of this for neuropathies resulting from metabolic disorders, such as diabetes, this also suggests that neurodegenerative disorders involving activation of these enzymes could have metabolic alterations as a part of their etiology, and not simply a downstream consequence of neurodegeneration and cell death. This prompts future work examining metabolic shifts preceding onset neurodegenerative diseases, and validates current efforts to establish metabolic biomarkers for this purpose.

Our findings build on previous mechanistic findings regarding Wnd/DLK in the regulation of axon degeneration, regeneration, and protection. In neurons, Wnd/DLK is activated by cytoskeletal disruption, phosphorylated by the cyclic adenosine monophosphate (cAMP) effector Protein Kinase A (PKA), and is restrained through turnover by the E3 ubiquitin ligase Hiw [12,135,136]. Wnd/DLK has also been linked to metabolism through the regulation of mitochondria [15,16,137]. In axon degeneration, Wnd/DLK regulates mitochondrial fission and works in parallel with mitochondrial dysfunction to decrease levels NMNAT2 and the microtubule regulator superior cervical ganglion-10/stathmin-2 (SCG10/STMN2) [15,16], with both NMNAT2 and SCG10/STMN2 being extensively-studied axon integrity factors important for both maintenance and regeneration [14,63,67,99,138–144]. In axon regeneration, Wnd/DLK regulates the localization of mitochondria to the regenerating axon [137]. External trophic supply through myelination is a core feature of peripheral axon support and recovery [145], and glial responses to injured neurons are regulated by Wnd/DLK [146–148]. For example, recent work by Duncan and colleagues found that neural apoptosis and axon loss following failure of remyelination is dependent on DLK signaling [148]. These studies link Wnd/DLK, metabolic support, and neurite integrity in disease and injury models. Our study demonstrates that disruption of metabolism through targeting of PyK in neurons is sufficient to activate both protective and degenerative Wnd/DLK signaling and downstream genetic changes. Future work examining how these functions can be uncoupled, such as inhibiting the neurodegenerative signal while enhancing the protective signal, could offer therapeutic strategies.

### Nuanced control of axon survival

One of the central aspects of neurodegeneration is stress response—the innate signaling activated in neurons entering axonal, synaptic, and/or cell body destruction. Over a century of research has made clear that neural stress responses can promote or inhibit neurodegeneration [33,149,150]. One of the most persistent questions centers on how mediators of stress response are able to regulate multiple, often conflicting, protective and destructive outcomes. Wnd/DLK is a prime example of this, as homologs of this protein have been linked to conserved mechanisms of axon degeneration, regeneration, and protection across disease models and species [33]. The biological basis for having disparate, seemingly conflicting mechanisms controlled by a single actor remains highly studied and extremely relevant for clinical care. In our study, we found that loss of PyK induced axonal and synaptic degeneration through Wnd/DLK. Given the known protective effects of Wnd/DLK following a conditioning lesion [58,96], and our JNK reporter data indicating Wnd/DLK genetic regulation in PyK-deficient motor neurons, we used physical injury at an earlier developmental stage to test for evidence of a protective response to PyK-deficiency. The delay in axon degeneration we observed post-injury supported this hypothesis, as did the partial dependence of this delay on Wnd/DLK signaling. These data in combination with our finding that Wnd/DLK promotes the progressive synaptic degeneration caused by PyK loss could indicate a rheostat system responsive to metabolic perturbations. This system likely converges on regulation of SARM1, which our experiments implicate as an effector in both the degenerative and protective responses to PyK loss. Knockdown of dSarm/SARM1 suppresses the synaptic fragmentation resulting from PyK-deficiency to a similar extent as knockdown of Wnd/DLK. Additionally, early 3rd instar larval motor axons show a significant reduction in expressed GFP-tagged dSarm/SARM1 localizing to the axon following PyK knockdown. Overexpression of dSarm/SARM1 in these axons fully abolishes the delay in injury-induced axon degeneration, supporting the hypothesis that this delay is due to reduced axonal dSarm/SARM1. Together, these data indicate that regulation of dSarm/SARM1 is central to both the neurodegenerative and neuroprotective responses to PyK loss in motor neurons, with Wnd/DLK as a potential upstream regulator (**Figure 7**). The biological purpose for having this type of rheostat could be in the conservation of resources by switching from preserving to sacrificing a neuron that has passed some established point of survival and function, preventing it from remaining a non-functioning metabolic burden.

**Figure 7:**
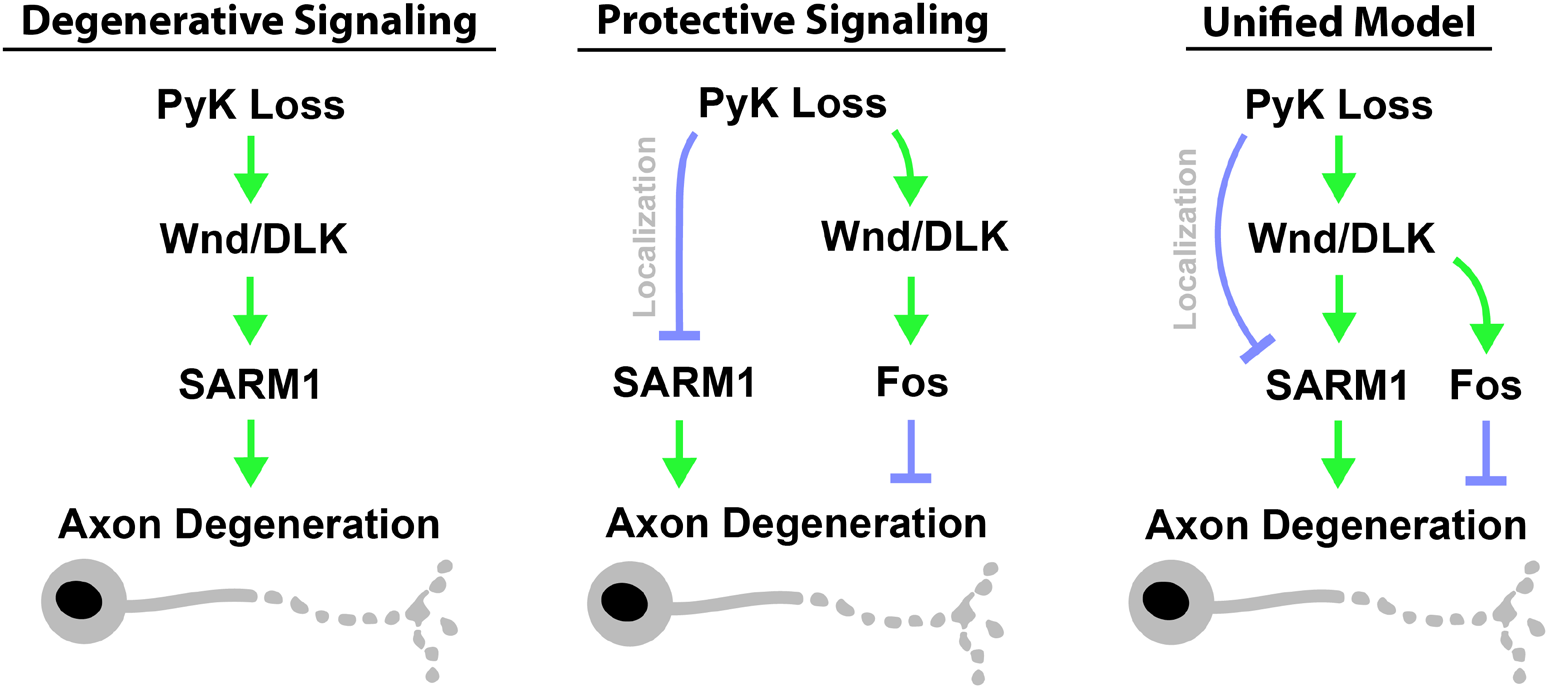
Model. We propose a model of two distinct signaling responses to metabolic disruption following PyK loss. The first pathway acts through Wnd/DLK and SARM1 to drive axonal and synaptic degeneration of PyK-deficient motor neurons. Prior to this degeneration, a protective response is dominant, as revealed by delayed degeneration of PyK-deficient axons cut from the neuron cell body earlier in larval development. This protective response reduces localization of SARM1 to the axon, which inhibits degeneration. Inhibition of Wnd/DLK significantly, but only partially, blocks this protective effect, suggesting that additional mechanisms may be acting to inhibit degeneration in response to PyK loss, potentially converging on regulation of SARM1 localization. Additionally, the partial dependence of this protection of Fos implicates a protective genetic program. Together, propose that these two responses to metabolic disruption act as a switch, suppressing degeneration in response to mild or short-term stress and promoting degeneration in response to high or chronic stress.

### Requirement for glycolysis and mitochondrial metabolism in neurons

One of the greatest questions in both basic and health neurosciences is the metabolic homeostasis of neurons, which is vital for understanding neural function and disease. Research on this question has been an especially vigorous regarding the dependence of neurons on glia for nutrient processing and supply [84–89]. Recent work has produced multiple perspectives across model systems, and neuron and glia subtypes, showing that neurons do or do not depend on glia for varies metabolites, including precursors to pyruvate [37,84,91,151–153]. In this study, we found that motor neuron-specific depletion of PyK causes progressive axon and synapse loss. This supports the hypothesis that some neuron types are dependent on metabolic flux through PyK, rather than relying solely on pyruvate equivalents from glia that could bypass PyK for entry into mitochondrial metabolism. Given that a major phenotype of this dependence is shown as synaptic loss and fragmentation of the axon, it is possibly that axonal metabolism specifically may depend on PyK. In context of the known importance of PyK for carbon input into the mitochondrial citric acid cycle, and the role of mitochondria in the maintenance and function of axons and synapses, our data could indicate that the mechanism of PyK-deficient axon degeneration involves axonal mitochondrial defects acting on Wnd/DLK and SARM1 to initiate the neurodegeneration process. In addition to the value of this information for the basic science of axonal metabolic regulation, it also highlights the potential benefits of targeting metabolism through diet or pharmaceutical methods in the treatment or prevention of neurodegenerative diseases.

## Limitations

This study approached the question of neural metabolism genetically, relying on RNAi-facilitated knockdown or Cas9-mediated knockout to target proteins and pathways. While this allowed for highly controlled cell-specific, consistent manipulations and analysis of outcomes in a physiological system, it did not directly examine metabolism in the neurons being studied. However, a recent study examining *pyk* mutant Drosophila larvae showed that disruption of this gene leads to metabolic changes also observed in human PyK deficiencies, including increases in glycolytic intermediates such as 2,3 bisphosphoglycerate (2,3-BGP) [36,154–157]. Additionally, as many of the manipulations relied on RNAi knockdowns and produced partial rescue phenotypes, we cannot conclude that Wnd/DLK and AP-1 are uniquely involved, but our experiments do support a necessary and robust role for them in context-specific responses to metabolic perturbations. Additional factors are likely involved and identifying them will be the focus of future work.

### Future directions and therapeutic implications

One of the surprising findings in this work was the discovery of an innate neuroprotective pathway activated by metabolic perturbation and regulated by the Wnd/DLK kinase. Given the prevalence of metabolic perturbations across neurodegenerative disorders, the existence of innate neuroprotective responses to these disruptions is highly relevant to development of therapeutic interventions, both as potential targets to enhance and as existing integrity mechanisms to avoid disrupting. For example, perturbations in glucose metabolism have been linked to amyotrophic lateral sclerosis (ALS) across a range of human and animal studies [5,34,116,158–163]. Both SARM1 and Wnd/DLK have also been implicated in ALS through genetic and mechanistic studies [26,122–125]. Recently, the first Phase I human clinical trial examining the GDC-0134 DLK inhibitor in patients with ALS was conducted [164]. This study was stopped due to elevated neurofilament light chain (NFL) levels in patient plasma, a biomarker of neurodegeneration [165], raising concens on the potential harmful effects of DLK inhibition [164]. This highlights the necessity for fully understanding the dual neuroprotective and neurodegenerative mechanisms of DLK so that its harmful activity can be specifically targeted without disrupting its protective effects. Therefore, studies that identify mechanisms and effectors specific to each are paramount.

Given our discovery of a genetic program activated in PyK-deficient motor neurons and regulated by Wnd/DLK, further investigations into the specific genes included in this program could uncover downstream effectors of Wnd/DLK signaling and provide an inroad for identifying targets for separating the neurodegenerative and neuroprotective functions of this pivotal enzyme.

## Acknowledgments

We thank Marc Freeman (Vollum Institute) for sharing the dSarm-GFP flies and Gregory Macleod (Florida Atlantic University) for sharing the pHusion-Ras flies. Fly stocks obtained from Bloomington Drosophila Stock Center (BDSC, NIH P40OD018537) and Vienna Drosophila Resource Center (VDRC, [166]) were used in this study. This research was funded by the National Institutes of Health (5T32DC000011-44 training grant to T.J.W., R01 NS069844/NS/NINDS to C.A.C., and NIH R01DK130875 to M.D.). This research was also funded by the Rita Allen Foundation, Klingenstein Fellowship in the Neurosciences, and NSF CAREER 1941822 (to M.D.).

## Supplemental Figures

**Supplemental Figure 1:**
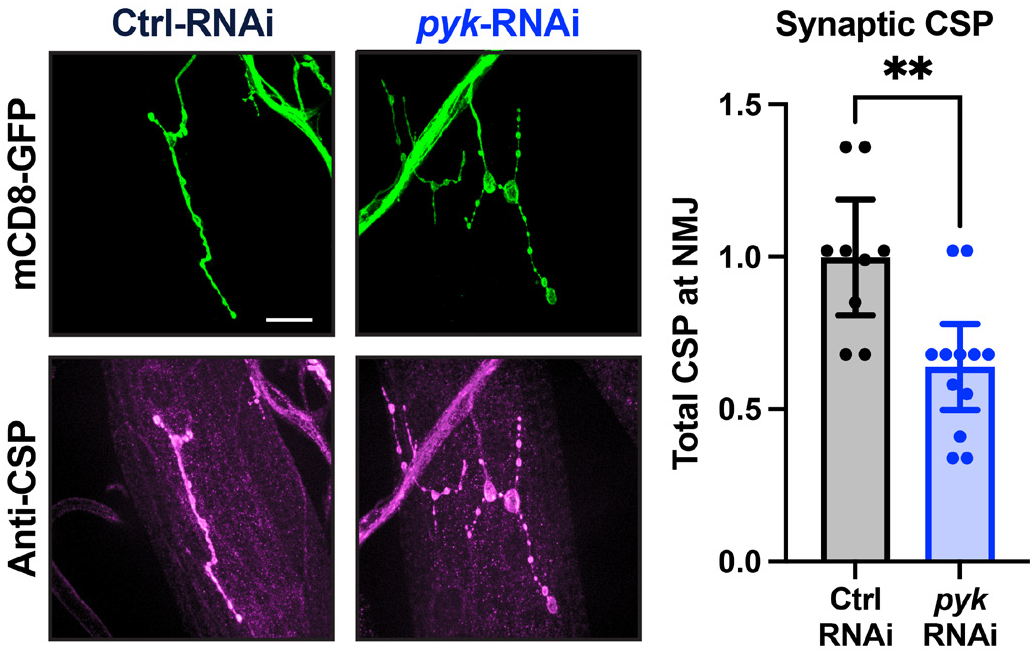
PyK knockdown alters NMJ structure. Neuromuscular junctions (NMJs) of early 3^rd^ instar larvae expressing control (Luciferase) RNAi or PyK RNAi via the pan-motor neuron driver D42-Gal4. Neuronal membrane visualized by expressed mCD8-GFP (green) with synaptic vesicles labeled by staining against cysteine string protein (CSP, magenta). Between 1-3 Muscle 4 NMJs per animal (innervating body segments 3-5) from 5 larvae per condition were quantified for total CSP staining and plotted individually. Statistical comparison is an unpaired t-test. Scale bar is 20 μm. Significance symbol: ** p<0.01.

**Supplemental Figure 2:**
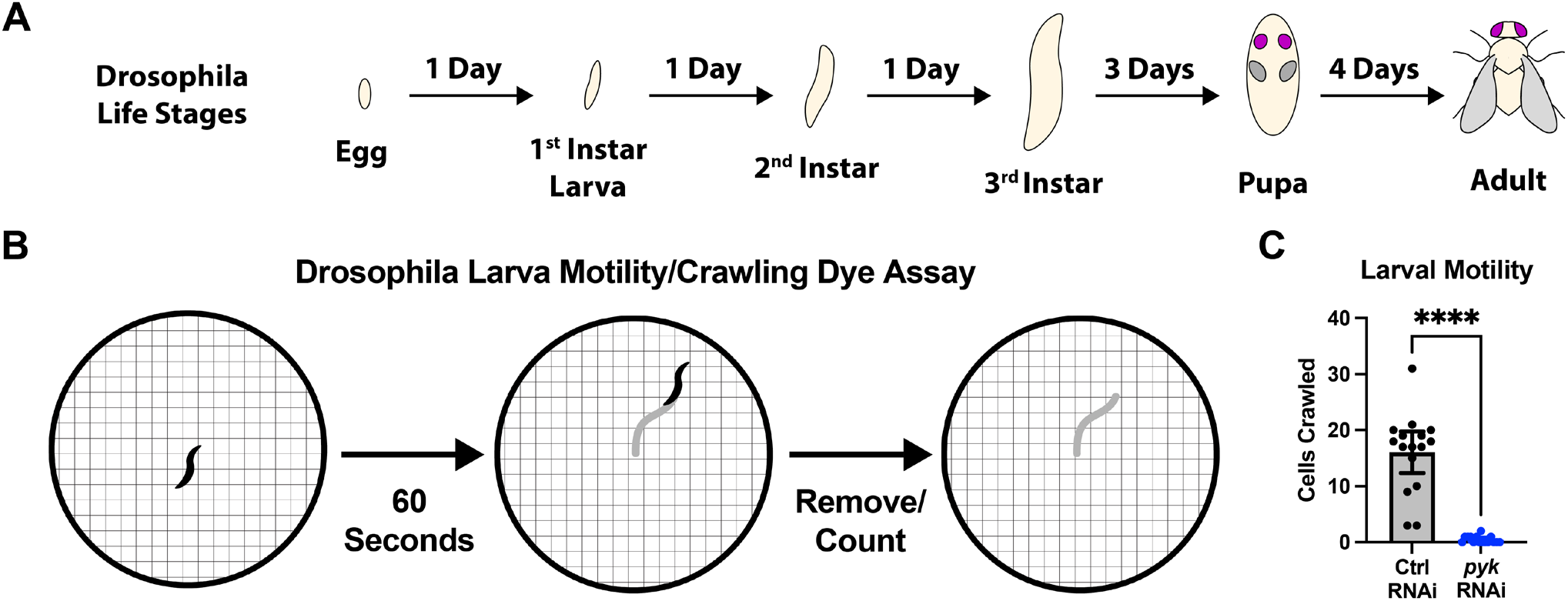
PyK deficiency in motor neurons causes motility defect in Drosophila larvae. **A)** Drosophila life stages with rough time durations. **B)** Schematic of larval crawling assay for measuring motility, described in methods. Larvae are placed in a drop of dye and allowed to crawl on a 10cm petri dish for 60s. The total number of 2mm cells marked with dye is counted. **C)** Cells crawled by late 3^rd^ instar larvae expressing control (Luciferase) RNAi or PyK RNAi in motor neurons (via D42-Gal4). 15-16 larvae per condition were tested with the larval crawling assay. Total cells crawled per larva are plotted individually. Statistical comparison is unpaired t-test. Significance symbol: **** p<0.0001.

**Supplemental Figure 3:**
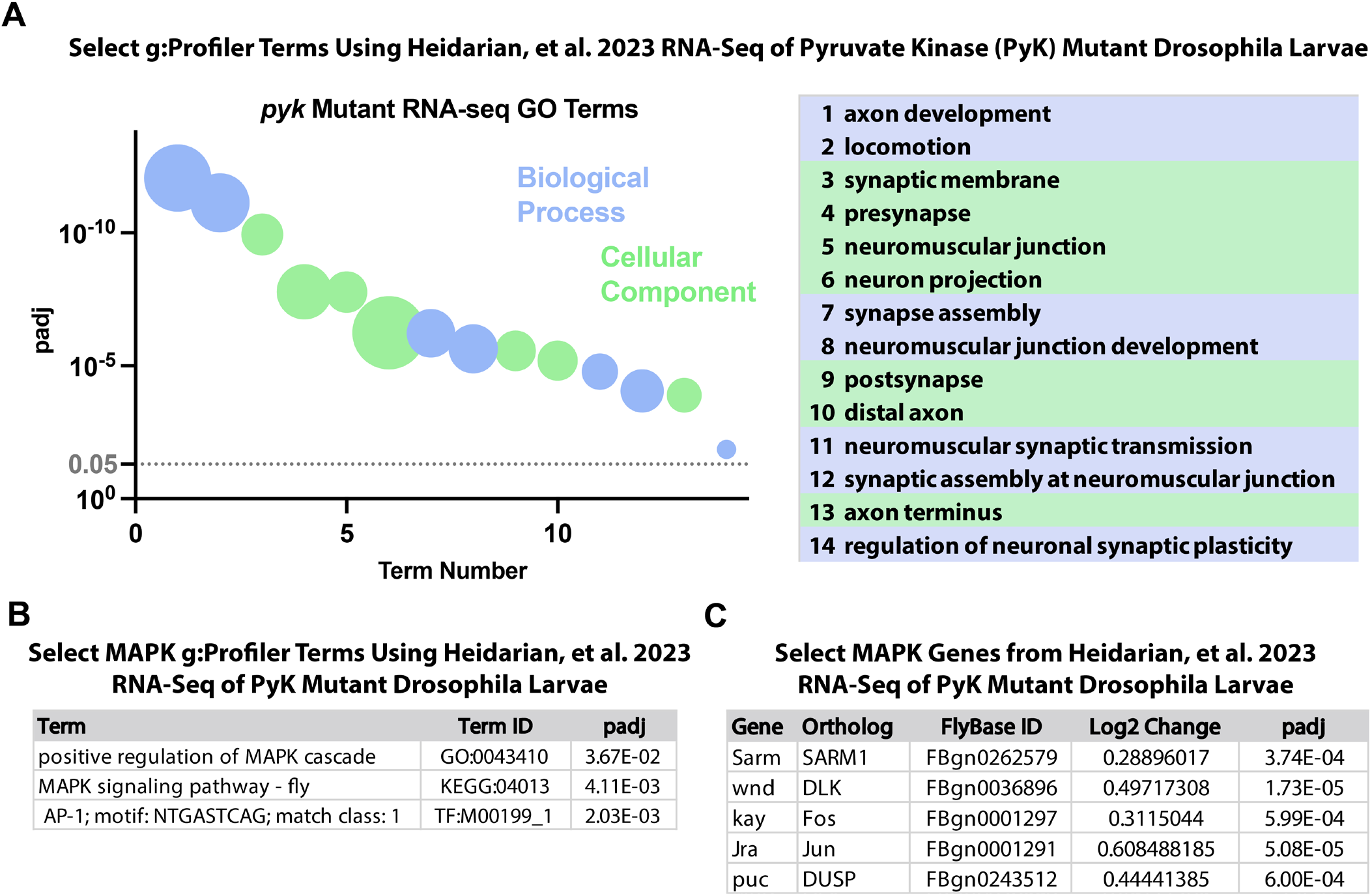
g:Profiler GO term analysis of Heidarian et al., 2023 RNA-seq of PyK mutant larvae. **A)** Select GO terms and significances using g.Profiler [47] enrichment analysis of significantly upregulated genes (padj<0.05) in PyK mutant larvae (PyK^23/31^) compared to heterozygote controls (PyK^23/+^) from Heidarian et al., 2023 [36]. The area of plotted dots shows the relative number of gene intersections for that term (values range from 10 to 164 genes). **B)** Select MAPK GO terms and significances using g.Profiler [47] enrichment analysis of significantly upregulated genes (padj<0.05) in PyK mutant larvae (PyK^23/31^) compared to heterozygote controls (PyK^23/+^) from Heidarian et al., 2023 [36]. **C)** Select MAPK gene Log2-fold changes and significances of PyK mutant larvae (PyK^23/31^) compared to heterozygote controls (PyK^23/+^) from Heidarian et al., 2023 [36].

**Supplemental Figure 4:**
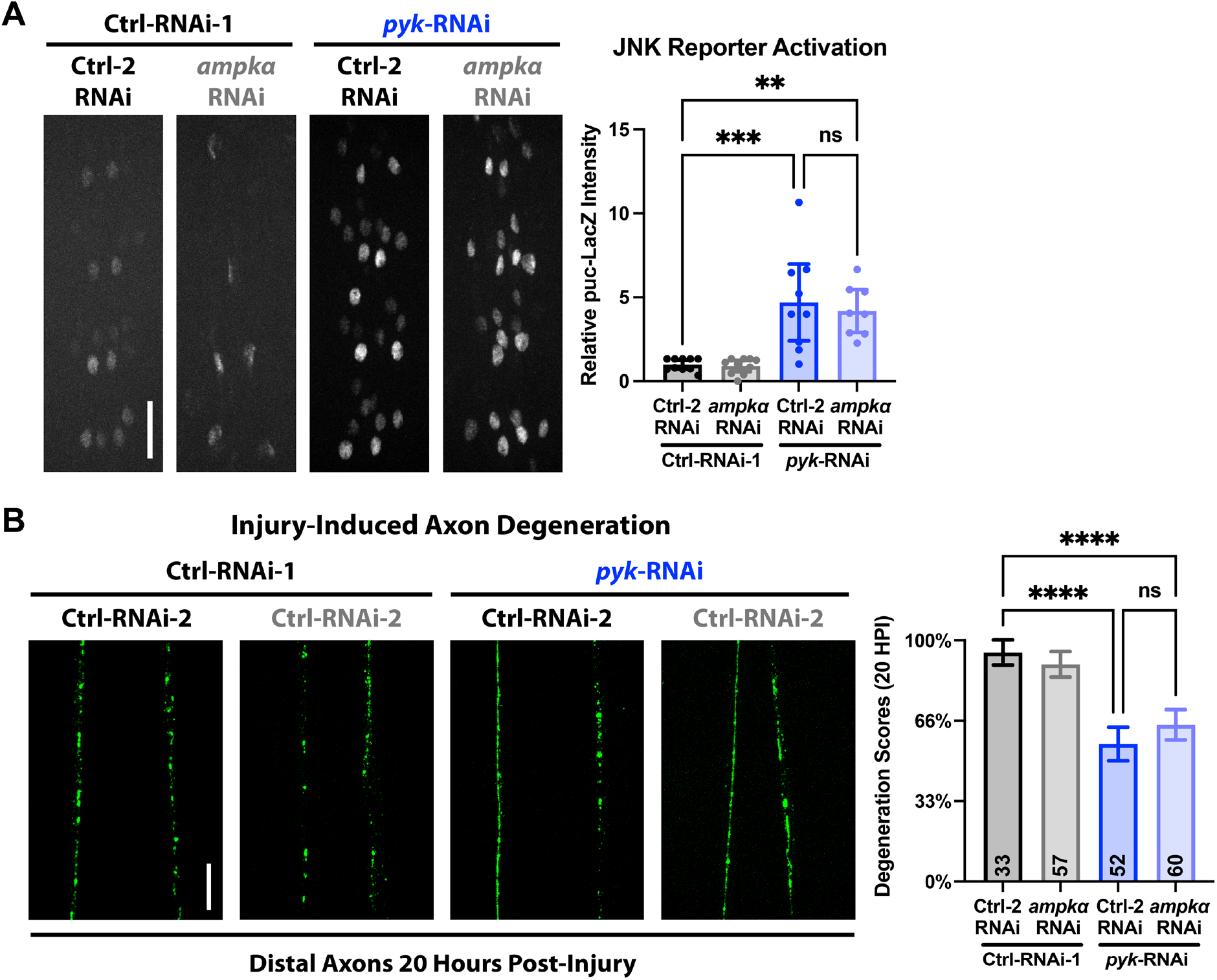
Co-knockdown of AMPKα does not suppress JNK signaling activation or delayed axon degeneration post-injury of PyK RNAi. **A)** Levels of β-Gal in the midline motor neuron nuclei of *puc*-LacZ mid 3^rd^ instar larvae expressing either *luciferase*-RNAi or *pyk*-RNAi in combination with either *lexA*-RNAi or *ampkα*-RNAi. Nuclei from motor neurons innervating body segments 4-7 (8-10 cells per corresponding segment, 32-40 total per animal) from 8-10 larvae per condition were quantified for β-Gal staining intensity and were plotted as animal averages. **B)** Distal injured motor neuron axon pairs of mid 3^rd^ instar larvae expressing *luciferase*-RNAi or *pyk*-RNAi in combination with either *lexA*-RNAi or *ampkα*-RNAi, with mCD8-GFP for visualization. Between 1-9 severed axon pairs each from 6-9 larvae per condition were scored blinded for extent of degeneration using a 4-point scale (0% - completely continuous, 33% - continuous with varicosities, 66% - partially continuous and partially fragmented, 100% - fully fragmented). Total number of axons pairs scored per condition shown on graph. All statistical comparisons are one-way ANOVA with Tukey correction for multiple comparisons. Scale bar is 20 μm. Significance symbols: * p<0.05, ** p<0.01, *** p<0.001, **** p<0.0001, ns = not significant.

